# Simultaneous quantification of ankle, muscle, and tendon impedance in humans

**DOI:** 10.1101/2021.12.08.471793

**Authors:** Kristen L. Jakubowski, Daniel Ludvig, Daniel Bujnowski, Sabrina S.M. Lee, Eric J. Perreault

## Abstract

**Objective:** Regulating the impedance of our joints is essential for the effective control of posture and movement. The impedance of a joint is governed mainly by the mechanical properties of the muscle-tendon units spanning it. Many studies have quantified the net impedance of joints but not the specific contributions from the muscles and tendons. The inability to quantify both muscle and tendon impedance limits the ability to determine the causes underlying altered movement control associated with aging, neuromuscular injury, and other conditions that have different effects on muscle and tendon properties. Therefore, we developed a technique to quantify joint, muscle, and tendon impedance simultaneously and evaluated this technique at the human ankle.

**Methods:** We used a single degree of freedom actuator to deliver pseudorandom rotations to the ankle while measuring the corresponding torques. We simultaneously measured the displacement of the medial gastrocnemius muscle-tendon junction with B-mode ultrasound. From these experimental measurements, we were able to estimate ankle, muscle, and tendon impedance using non-parametric system identification.

**Results:** We validated our estimates by comparing them to previously reported muscle and tendon stiffness, the position-dependent component of impedance, to demonstrate that our technique generates reliable estimates of these properties.

**Conclusion:** Our approach can be used to clarify the respective contributions from the muscle and tendon to the net mechanics of a joint.

**Significance:** This is a critical step forward in the ultimate goal of understanding how muscles and tendons govern ankle impedance during posture and movement.

## I. Introduction

The mechanics of a joint arise, in part, from the mechanical properties of the muscles and tendons that span it [3]. Therefore, the mechanical properties of our muscles and tendons have a significant impact on how we interact with our physical environment. Despite this, our current understanding of how joint mechanics are varied to enable our seamless interactions is almost entirely from measurements of joints and whole limbs, which provide little insight into the mechanics of the muscle and tendon. The contributions from muscle and tendon to the mechanics of a joint have also been approximated using musculotendon models [5–7]. However, the lack of measurements of both muscle and tendon mechanics is a critical limitation as muscle and tendon mechanical properties can be altered differently by aging, pathology, or injury. Thus, distinguishing how changes within the muscle and tendon contribute to the impaired control of posture and movement is crucial for rehabilitation and targeted treatments.

Currently, there are no methods that can simultaneously quantify the contributions from the muscle and tendon to the impedance of a joint across all levels of muscle activation. Impedance quantitatively describes the mechanical properties of a joint as the dynamic relationship between an imposed displacement and the resultant torque [3]. While joint impedance has been quantified under a variety of conditions [9–12], these measures cannot distinguish the contributions from the muscles and tendons to the net mechanics of the joint. Others have quantified muscle and tendon stiffness, the static component of impedance, but with several limitations. First, a majority of the methods used to date can assess the stiffness of either the muscle or tendon, not both simultaneously [8, 13, 14]. This limits the understanding of the interaction between the muscle and tendon and how they contribute to the net mechanics of the joint. Second, it is commonly assumed that the stiffness of the tendon is constant [8, 13, 14]. This is despite evidence that it varies non-linearly with load, increasing at low strains before reaching a constant value at high strains [15]. The assumption of tendon stiffness being constant has been justified by restricting estimates to higher contraction levels, typically above ~30% maximum voluntary contraction (MVC) [16]. While these estimates have provided valuable information, they are not directly relevant to many essential tasks such as walking and standing, where the mean muscle activation remains below 30% MVC [17]. One study has assessed muscle and tendon stiffness at a lower level of muscle activation [18]. However, it relied on a Hill-type muscle model to calculate the ratio of muscle stiffness to tendon stiffness. Any errors in the model would impact their results, and Hill-type models are known to have limited accuracy, especially during dynamic conditions [19]. Due to these limitations, it remains unclear how the muscle and tendon contribute to the impedance of the joint.

The objective of this work was to develop a quantitative *in vivo* technique for simultaneous estimation of human joint, muscle, and tendon impedance. We evaluated our technique at the human ankle, though it could be applicable to other joints provided that the required measurements are feasible. The experimental approach leveraged controlled displacements of the ankle, measures of ankle torque, and motion of the medial gastrocnemius muscle-tendon junction (MTJ) obtained from B-mode ultrasound. Our analysis of these data relied on nonparametric system identification, as has previously been employed to estimate the impedance of individual joints [10, 12, 20]. The measures of muscle-tendon motion that we collected allowed us to estimate muscle and tendon impedance in addition to joint impedance. We validated our experimental estimates against scaled measurements from animal and cadaveric studies as well as an *in vivo* study on the Achilles tendon that made measurements below 30% MVC [1, 4, 8]. Our findings demonstrate the utility of this technique to obtain reliable estimates. Our method enables future work that can focus on determining how muscle and tendon contribute to the impedance of a joint over a range of conditions. Part of this work has been presented previously in abstract form [21].

## II. Methods

The main contributions of this manuscript are the experimental paradigm and analysis methods that we developed to quantify joint, muscle, and tendon impedance across a range of activation levels. First, we describe the analysis methods and their assumptions. We then present our experimental methods demonstrating the use of this technique at the human ankle. We expect that the same methodology should work for other joints at which the necessary experimental measures can be made. The key experimental measures are ankle angle, ankle torque, and displacement of the MTJ (Fig 1a) (see *Experimental Protocol* for details on how we collected these measures).

**Fig 1.**
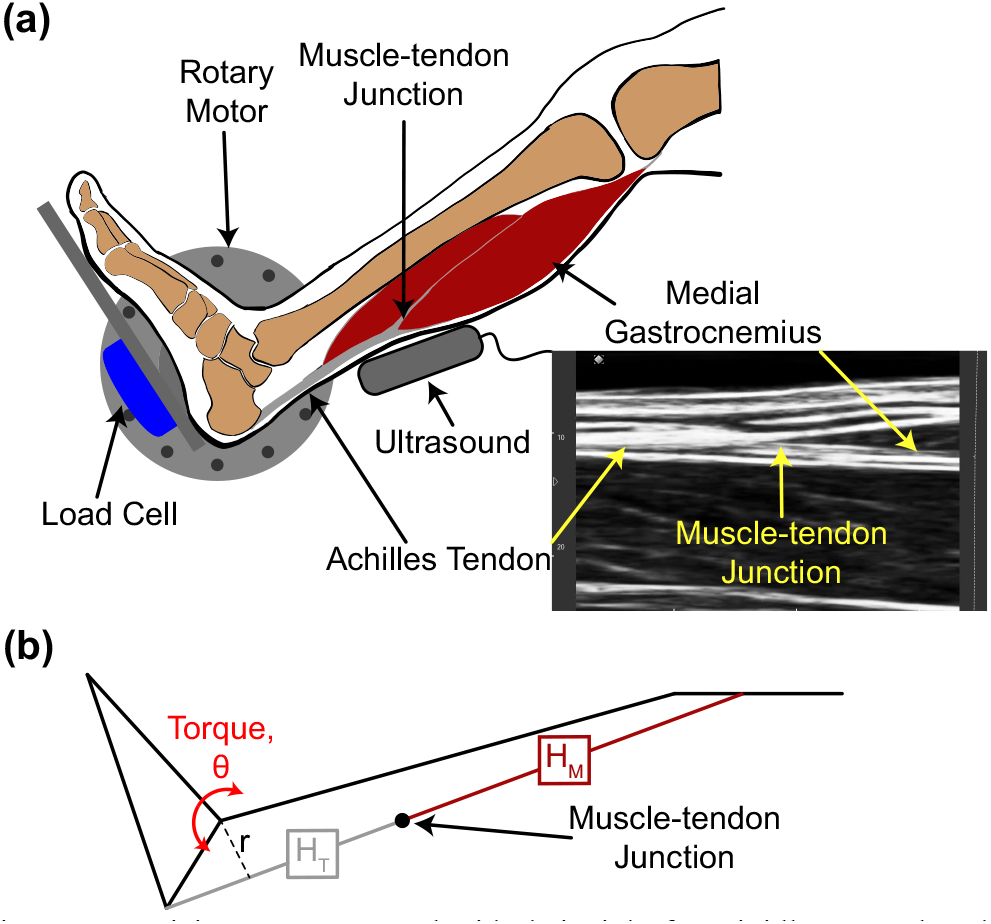
(a) Participants were seated with their right foot rigidly secured to the rotary motor. The rotary motor precisely controlled the participant’s ankle angle while the load cell measured the resultant ankle torque. An ultrasound system (B-mode) imaged the muscle-tendon junction of the medial gastrocnemius. The images were manually digitized to quantify muscletendon junction displacement. (b) We modeled the muscle (*H_M_*) and tendon (*H_T_*) as two impedances in series, connected to the foot with a known moment arm (*r*). We used ankle angle (*θ*), torque, and muscle-tendon junction displacement to calculate ankle, muscle, and tendon impedance.

### A. Estimates of ankle, muscle, and tendon impedance

We performed a non-parametric system identification analysis on our experimental measures to estimate the dynamics of the ankle, muscle, and tendon [22]. A critical component of this analysis is using a noise-free input [3]. Thus, we performed our analysis such that ankle angle, which is precisely controlled with a rigid manipulator, is always treated as the input. Our analysis is a small signal analysis that assumes our experimental variables are small perturbations around an operating point.

First, ankle impedance (*H_A_*) was estimated as the relationship between an imposed ankle rotation (*θ*) and the torque generated in response (1; Fig 2) [3, 9, 10]

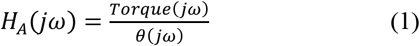

where 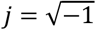 and *ω* is the frequency in radians per second.

**Fig 2.**
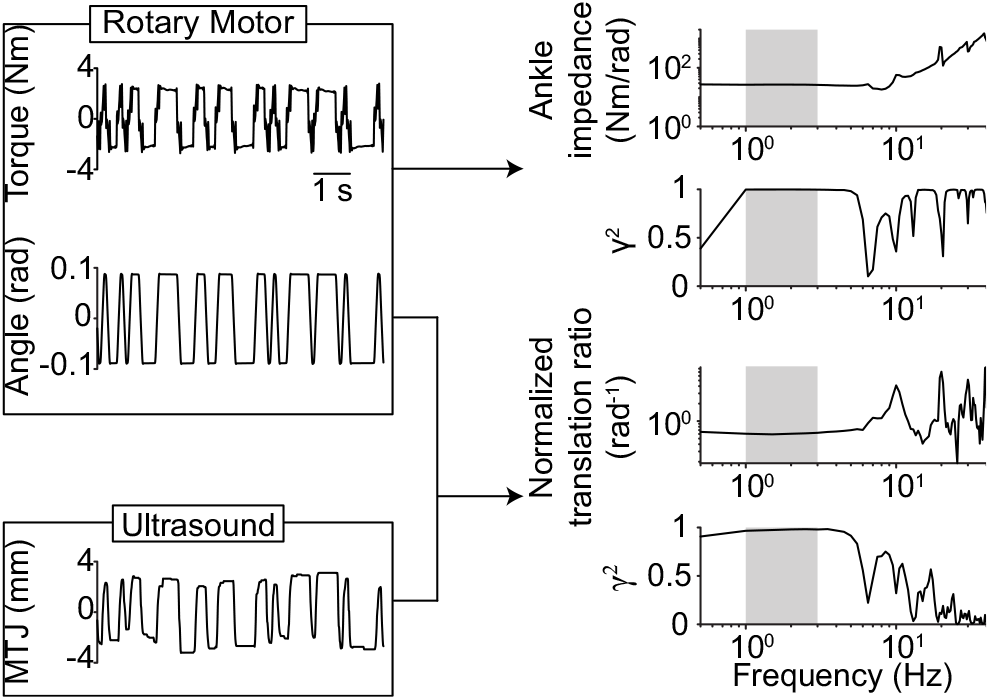
Representative data used to estimate ankle impedance and the normalized translation ratio. We used ankle angle and ankle torque data to calculate the frequency response function quantifying ankle impedance and the corresponding coherence (γ^2^). We used ankle angle and muscle-tendon junction (MTJ) displacement to calculate the frequency response function quantifying the normalized translation ratio. The MTJ displacement is from the medial gastrocnemius. Values between 1 to 3 Hz used to compute stiffness and the mean coherence are shaded.

Next, we estimated the relationship between an imposed ankle rotation and the resultant displacement of the MTJ, which we refer to as the *translation ratio T_R_* (2) (Fig 2).

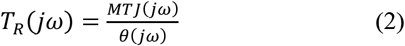

*H_A_* and *T_R_* can be used to estimate muscle (*H_M_*) and tendon (*H_T_*) impedance under the assumption that the muscle and tendon are connected in series [23], rigidly attached to the calcaneus (Fig 1b), and that the proximal end of the muscletendon unit is fixed. With these assumptions, the displacement of the MTJ is given by:

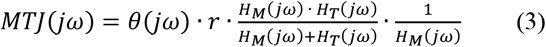

where *r* represents the Achilles tendon moment arm. As such, *T_R_* represents, when scaled by the moment arm, the ratio of net musculotendon impedance to the impedance of the muscle. We used a single Achilles tendon moment arm for all calculations since it does not scale with anthropometric data [24, 25]. An estimated value of 51.4 mm was used, which was the mean across participants from Clarke et al. [24] with an ankle angle of 90°. All results for *T_R_* are normalized by the moment arm estimate, which we referred to as the normalized translation ratio, 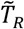.

Muscle impedance, by definition, is the relationship between changes in muscle length and the resultant changes in muscle force. When applying our technique to the human ankle, our algorithm assumes that *H_M_* represents the net impedance of the triceps surae and *H_T_* represents the net impedance of the Achilles tendon (Fig 1b; see *Results* and *Discussion* where we test the validity of this assumption). It also assumes that all measured plantarflexion torque about the ankle is fully transmitted through the Achilles tendon, allowing us to estimate muscle force by dividing ankle torque by the Achilles tendon moment arm. Thus, we can determine muscle impedance from (1) and (2).

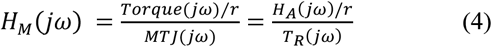

Tendon impedance can then be determined from (2) and (3).

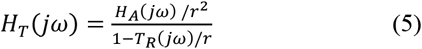

### B. Experimental set-up and protocol

To validate our method, we collected data from seven individuals (age = 26 ± 3 years, (mean ± std); height = 1.76 ± 0.09 m; body mass = 68 ± 13 kg; 4 males). All participants were right leg dominant and had no history of neuromuscular or musculoskeletal injuries to their right leg. Participants provided informed consent prior to participation. The Northwestern University Institutional Review Board approved this study (STU00009204 & STU00213839).

Participants were seated in an adjustable chair (Biodex Medical Systems, Inc. Shirley, NY, USA). The knee was stabilized at 15° of flexion with a brace (Innovator DLX, Ossur, Reykjavik, Iceland), which prevented movement at the proximal end of the biarticular medial and lateral gastrocnemii. We rigidly secured the participant’s foot to an electric rotary motor (BSM90N-3150AF, Baldor, Fort Smith, AR, USA) via a custom-made fiberglass cast. Ankle position was set at 90° of flexion. The rotation of the motor was restricted to the sagittal plane. We aligned the ankle center of rotation in the sagittal plane with the center of rotation of the motor. Electrical and mechanical safety stops limited the rotation of the motor within the participant’s range of motion. An encoder integrated within the motor measured ankle angle, while a six-degree-of-freedom load cell (45E15A4, JR3, Woodland, CA, USA) measured all ankle forces and torques. The analog data were passed through an antialiasing filter (500 Hz using a 5-pole Bessel filter) and sampled at 2.5 kHz (PCI-DAS1602/16, Measurement Computing, Norton, MA, USA). Ankle angle was simultaneously recorded using a 24-bit quadrature encoder card (PCI-QUAD04, Measurement Computing, Norton, MA, USA).

Electromyography (EMG) data were collected from the medial and lateral gastrocnemii, soleus, and tibialis anterior using single differential bipolar surface electrodes (Bagnoli, Delsys Inc, Boston, MA, 10 mm interelectrode distance). Standard skin preparation techniques were used before applying each electrode to the skin [26]. Electrodes were placed on the belly of the respective muscle. EMG data were collected for the visual feedback provided to the subjects.

We rigidly secured a B-mode ultrasound probe (LV7.5/60/128Z-2 Telemed, Lithuania) to the leg to image the medial gastrocnemius MTJ. We positioned the probe parallel to the muscle belly (longitudinally) such that the MTJ was centered on the image. Ultrasound data had a mean frame rate of 124 Hz and were synchronized with all measurements from the rotary motor. We saved all images as video files for further analysis.

Our analysis assumes that we are estimating the net impedance of the triceps surae and Achilles tendon. This implies that MTJ displacement, the subsequent translation ratio, and muscle and tendon impedance estimates are matched across the three muscles of the triceps surae. To evaluate the validity of this assumption, we made measurements while imaging the MTJ of the medial and lateral gastrocnemii and soleus in three of the seven subjects. For each muscle, we positioned the probe parallel to the muscle such that the MTJ was centered on the image. Due to time restrictions, only two MTJ’s could be imaged per day. Therefore, we imaged the medial and lateral gastrocnemii MTJ’s on Day 1 and the medial gastrocnemius and soleus MTJ’s on Day 2. Due to a global pandemic, Day 1 and Day 2 testing was separated by an average of 490 days. On Day 1, the ultrasound data had a mean frame rate of 124 Hz. Due to its structure and location, when imaging the soleus MTJ, we altered the ultrasound scanning parameters, decreasing the mean frame rate to 62 Hz. Day 2 images of the medial gastrocnemius MTJ were collected using the same parameters as for the soleus. On both days, muscle imaging order was randomized.

To quantify ankle, muscle, and tendon impedance, the rotary motor applied small rotational perturbations to the ankle while measuring the resultant ankle torque and displacement of the MTJ. Pseudorandom binary sequence (PRBS) perturbations with an amplitude of 0.175 radians, a maximum velocity of 1.75 radians per second, and a switching time of 153 ms were used. This profile is similar to what has previously worked well for estimating ankle impedance [27]. To determine the validity of our technique across a range of plantarflexion torques, participants produced varying levels of constant plantarflexion torque while the motor applied small perturbations about an otherwise isometric position. For each trial, we instructed participants to match a plantarflexion torque target with the aid of real-time visual feedback. We also provided real-time feedback of tibialis anterior EMG to prevent co-contraction. Adequate practice was provided for participants to become proficient with the visual feedback. We tested plantarflexion torque levels of 0% to 30% MVC in 5% increments. Participants completed three 65-second trials for each torque level. For Day 2 measurements, the trial duration was decreased to 55 seconds, which we had determined was sufficient for purposes of system identification. We randomized the trial order for each participant. Participants were allowed to rest at least 1 minute between trials to prevent fatigue. Longer rest breaks were provided after trials that required higher levels of muscle activation (>20% MVC). Participants were repeatedly asked if they were experiencing fatigue. If the participant indicated that they were fatigued, a rest break was provided until they indicated they were no longer fatigued.

### C. Data processing and analysis

We performed all data processing and analysis using custom-written software in MATLAB (Mathworks, Natick, MA, USA). We manually digitized the MTJ within each frame of the ultrasound videos. The ultrasound device has a variable interframe interval, which we accounted for during our processing [28]. The angle and torque data were collected at a different sampling rate from the ultrasound images. As such, ultrasound data were linearly interpolated to the higher sampling rate for all further analyses.

The measured ankle torque includes inertial and gravitational contributions from the mass properties of the apparatus rigidly connecting the foot to the motor. As such, we ran a single trial without the foot in place so that the torque from the apparatus could be estimated and removed from the net torque measured in each trial [29].

To characterize the ankle impedance and translation ratio, we computed the non-parametric frequency response functions between the controlled input and the measured output (Fig 2) [3]. The quality of the frequency response functions was assessed using input-output coherence (γ^2^; Fig 2) [3], and the percentage of variance accounted for (VAF) between the measured and modeled output variables. We computed muscle and tendon impedance from the estimated ankle impedance and translation ratio frequency response functions ((4) & (5)). Stiffness is the dominant contributor to impedance at low frequencies. As such, we approximated ankle, muscle, and tendon stiffness, and the normalized static translation ratio as the average magnitude of the corresponding frequency response functions over 1 to 3 Hz, where these quantities were nearly constant, indicating a static relationship between input and output (Fig 2, shaded region). Coherence was also high in this region, indicating that there was a linear relationship between the measured inputs and outputs in this region. The VAF assessed how well a linear model fits the data overall, while the averaged coherence between 1-3 Hz quantified the fit of the frequency response functions at the frequencies relevant to stiffness. We used these values to summarize our results across participants and tested conditions.

### D. Comparison with previous literature

We compared our muscle and tendon stiffness estimates to the literature since muscle and tendon impedance have not previously been quantified *in vivo* in humans. Muscle stiffness was compared to previous measurements of short-range stiffness in the feline hindlimb [1], scaled to the human triceps surae. This approach has been shown to work well for estimating muscle contributions to the net stiffness of the human ankle, knee, elbow, and shoulder [5–7]. We assumed that the stiffness of the three muscles within the triceps surae are in parallel and sum linearly. Muscle short-range stiffness depends on muscle force and optimal fascicle length, such that different levels of load-sharing between the triceps surae muscles could alter the net muscle stiffness. Therefore, to determine estimates of the triceps surae stiffness from the literature, we included variability in load-sharing between these muscles based on previously reported values [2]. Total plantarflexion force was distributed such that the lateral gastrocnemius had 6.0 ± 2.5%, the medial gastrocnemius had 26.2 ± 11.8%, while the soleus had 67.8 ± 11.6% of the total force [2]. The force distribution among the triceps surae muscles was based on the physiological cross-sectional area and the activity within individual muscles during a submaximal voluntary contraction [2]. We used optimal fascicle length values of 6.4 ± 0.7 cm, 5.4 ± 0.8 cm, and 3.9 ± 0.8 cm for the lateral gastrocnemius, the medial gastrocnemius, and the soleus, respectively [2]. These previously reported distributions were used to generate expected distributions for triceps surae short-range stiffness, to which our experimental results could be compared. The model of muscle short-range stiffness we employed [1] does not include a passive component. Therefore, we added the passive muscle stiffness estimated in our experimental values to our model-based estimates of muscle short-range stiffness for all comparisons.

To our knowledge, there is only one previous *in vivo* measurement of Achilles tendon stiffness at activation levels similar to those in our study [8]. Therefore, in addition to comparing our results to the *in vivo* study, we also compared our tendon stiffness estimates to scaled measurements from tensile testing on cadaveric samples [4]. To scale these measurements, we used previously reported *in vivo* measurements of Achilles tendon cross-sectional area (56 mm^2^) and resting length (252 mm) [30].

### E. Sensitivity analyses

Estimates of muscle and tendon stiffness are sensitive to the assumed value of the Achilles tendon moment arm [6]. We performed a sensitivity analysis to determine the magnitude of this effect. We estimated muscle and tendon stiffness across the tested range of plantarflexion torques allowing the error in the moment arm to vary from −13.6% to 13.6% of the assumed value, which corresponds to a 7 mm difference. This range is twice the standard deviation across the participants reported by Clarke et al. [24]. For this analysis, estimates of ankle stiffness and the normalized static translation ratio are needed. These values were obtained from the experimental data (see Fig 6c & d). We defined muscle and tendon sensitivity as the percent change in stiffness divided by the percent change in the moment arm over the tested range of values.

### F. Statistical Analysis

We sought to determine if our estimates of ankle, muscle, and tendon stiffness differed between days and when imaging the medial gastrocnemius, lateral gastrocnemius, and soleus. Since our muscle and tendon stiffness estimates are based on our ankle stiffness and normalized static translation ratio estimates, for this analysis, we only tested for differences in ankle stiffness and the normalized static translation ratio, our two dependent variables. Ankle stiffness was modeled by a linear mixed-effects model. A generalized linear mixed-effects model with a power link function with a value of −1.9 was used to characterize the normalized static translation ratio since this varied non-linearly with ankle torque. This link function was selected empirically and evaluated to ensure a goodness of fit and that the resulting model errors were normally distributed. For all models, subject was treated as a random factor and plantarflexion torque as a continuous factor. When comparing across days, day was treated as a fixed factor. When comparing across muscles, muscle (medial and lateral gastrocnemii and soleus) was treated as a fixed factor. We performed all statistical analyses in MATLAB. For all models, we used a restricted maximum likelihood method and Satterthwaite corrections [31]. Significance was set *a priori* at α = 0.05. R^2^ were computed after removing subject-specific intercepts. Additional information on the mixed-effects models used for all analyses can be found in the Supplemental Material. All values, unless otherwise noted, are reported as the mean ± standard deviation.

## III. Results

### A. Ankle impedance and translation ratio

We first present results on data collected while imaging the medial gastrocnemius MTJ. Justification for this choice and comparisons to the results obtained when imaging the soleus and lateral gastrocnemius MTJs are presented in Section III-C below.

The non-parametric frequency response functions used to characterize ankle impedance and the normalized translation ratio fit the data well. The ankle impedance estimates had a mean VAF of 91 ± 6%, while the mean VAF for the normalized translation ratio was 70 ± 20%. For both estimates, coherence was high from approximately 1 to 6.5 Hz (Fig 3b & f), indicating that a linear description of these relationships accounts for most measured data variance within this frequency range. The drop in coherence at 6.5 Hz was due to the applied perturbations having reduced power above 6.5 Hz (Fig 3b, red arrow).

**Fig 3.**
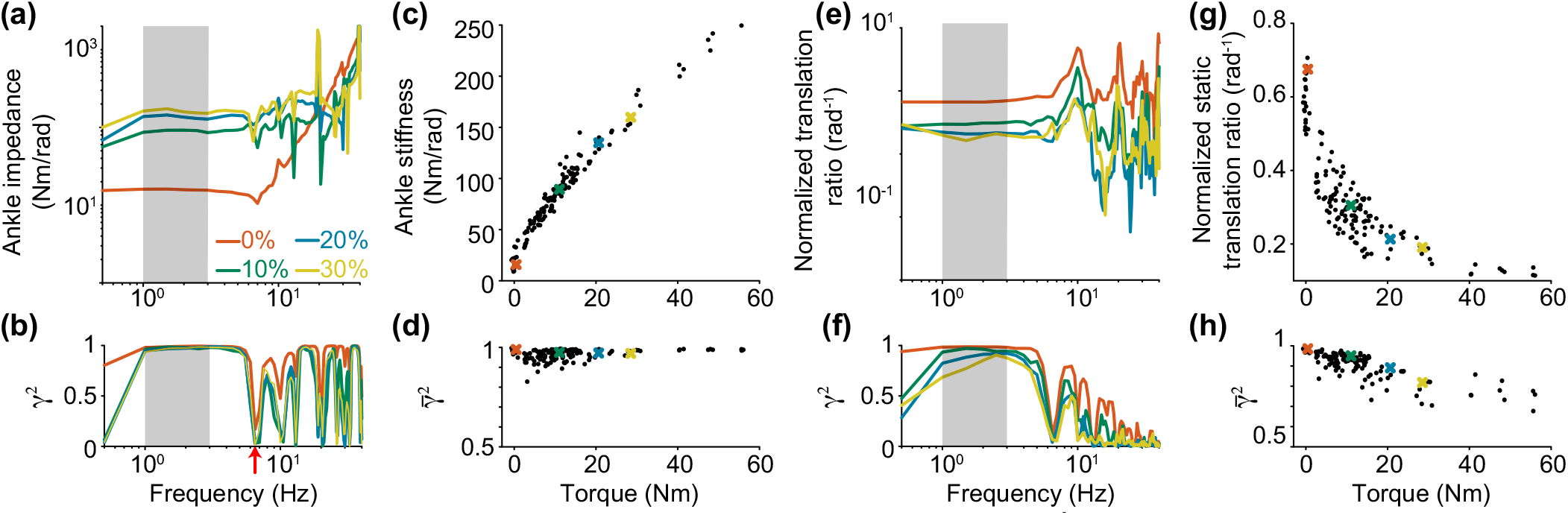
(a) Representative ankle impedance frequency response functions and (b) coherence functions (γ^2^). (e) Representative normalized translation ratio frequency response functions and (f) coherence functions (γ^2^). Different colors represent different levels of plantarflexion torque. Values between 1 to 3 Hz that were used to compute stiffness, normalized static translation ratio, and the corresponding mean coherences are shaded. (c) Ankle stiffness and (d) the corresponding average low-frequency coherence 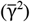 for all subjects. (g) The normalized static translation ratio and (h) the corresponding average low-frequency coherence 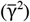 for all subjects. Each data point represents a trial. The colored x’s correspond to trials in a, b, e & f. The red arrow at 6.5 Hz denotes the drop in coherence due to the applied perturbations. having reduced power above 6.5 Hz.

The ankle impedance and normalized translation ratio varied as participants increased their voluntary torque (Fig 3a & e). The increase in ankle impedance with plantarflexion torque is consistent with previous results [32]. As plantarflexion torque increased, we also observed a decrease in the normalized translation ratio (Fig 3e), indicative of muscle impedance increasing more than tendon impedance ((2) & (3)).

Given that the magnitudes of the ankle impedance and normalized translation ratio were nearly constant from 1 to 3 Hz (Fig 3a & 3e), the magnitudes of these frequency response functions were averaged over this frequency range to obtain an estimate of the static system responses. The static component of impedance represents stiffness. Ankle stiffness increased with increasing torque, consistent with previous results (Fig 3c) [10]. As plantarflexion torque increased, the normalized static translation ratio decreased non-linearly, with the largest change coming at lower torque levels (Fig 3g).

Both ankle stiffness and the normalized static translation ratio had high average low-frequency coherence across all levels of plantarflexion torque (Fig 3d & h). The average low-frequency coherence was 0.97 ± 0.03 and 0.91 ± 0.07 for ankle stiffness and the normalized static translation ratio, respectively. There was a slight decrease in the average low-frequency coherence for the normalized static translation ratio as plantarflexion torque increased (Fig 3h). This was likely caused by a decrease in MTJ displacement as torque increased, decreasing the magnitude of the output signal.

Our values of ankle stiffness are similar to previously reported values. Kearney et al. [27] used slightly smaller perturbations (0.135 rad) and found that ankle stiffness varied from approximately 52 Nm/rad to 135 Nm/rad across their 5 participants when the plantarflexion torque was between 4-6 Nm. Our ankle stiffness values between 4-6 Nm were approximately 60 Nm/rad. Our values being on the lower end of the previously reported values could be due to slight differences in the participant’s knee angle and the tested perturbation amplitudes between the studies.

During all trials, the tibialis anterior remained quiet. The mean tibialis anterior activity was 1.1%. Therefore, tibialis anterior activity did not significantly contribute to our measurements.

### B. Muscle and tendon impedance

Both muscle and tendon impedance increased with an increase in plantarflexion torque (Fig 4a & b). The magnitude of these frequency response functions, computed from the estimated ankle impedance and normalized translation ratio, were also nearly constant from 1 to 3 Hz. Consequently, these were also summarized by their static values computed over these frequency ranges to provide estimates of muscle and tendon stiffness. We observed a linear increase in muscle stiffness as musculotendon force increased (Fig 5a). Tendon stiffness also increased, but non-linearly (Fig 5b).

**Fig 4.**
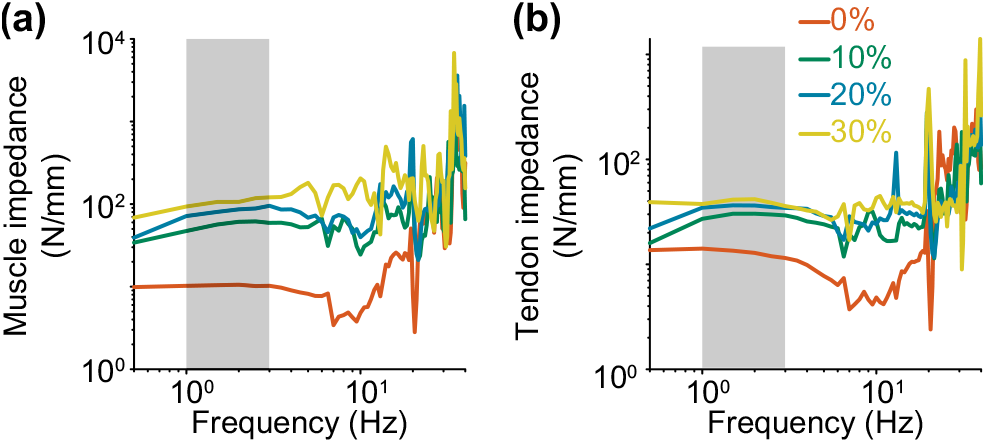
(a) Representative muscle impedance and (b) tendon impedance frequency response functions. Different colors represent different levels of plantarflexion torque. Values between 1 to 3 Hz that were used to compute stiffness are shaded.

**Fig 5.**
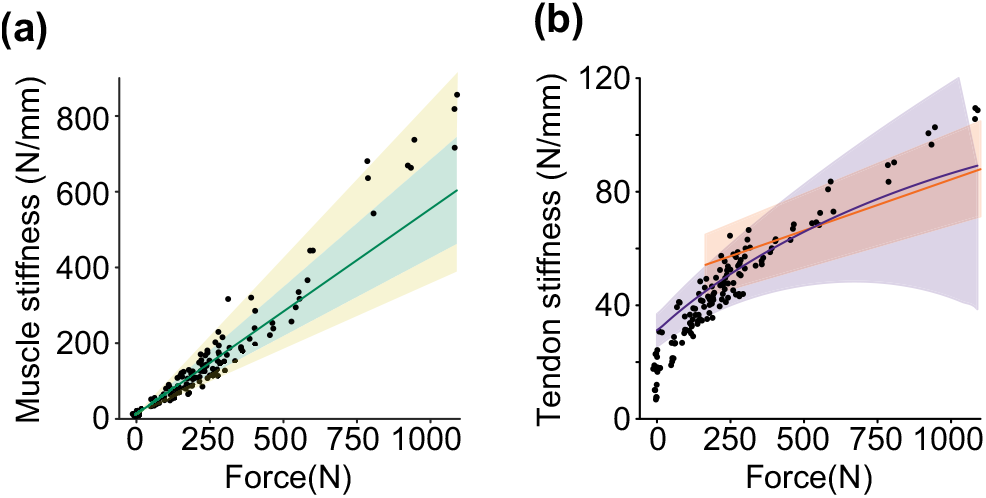
(a) Muscle stiffness estimates (black) and measurements of muscle short-range stiffness scaled to the geometry of the triceps surae (solid green line) [1]. We estimated muscle short-range stiffness with force scaled within each muscle of the triceps surae. The green shaded region tested for the variability in the force within each muscle, while the shaded yellow region tested for variability in the optimal fascicle length [2]. Nearly all of our estimates of muscle stiffness fall within scaled measurements of muscle shortrange stiffness. (b) Tendon stiffness estimates (black) and scaled measurements from Shaw and Lewis (purple) [4] and data adapted from Maganaris and Paul (orange) [8]. Nearly all of our estimates of tendon stiffness are within one standard deviation of the mean of the models estimated from the measures from Shaw and Lewis [4] and *in vivo* measurements from Maganaris and Paul [8] (shaded regions).

We compared our estimates of muscle stiffness to direct measurements of short-range stiffness from animals [1], scaled to the geometry of the triceps surae. Muscle short-range stiffness depends on the stress within a muscle and the length of its fibers. We, therefore, assessed how typical variation in load sharing between the muscles of the triceps surae (green) and optimal fascicle length (yellow) would influence the muscle stiffness-force relationship (Fig 5a) [2]. We found that our estimates of muscle stiffness are comparable to the previously published direct measurements from animals. Nearly all of our estimates of muscle stiffness are within the muscle short-range stiffness estimates that include physiological variability in force sharing and fascicle length within the triceps surae. For a majority of our estimates, including variability in force sharing was sufficient to account for the variability in our estimates.

We also compared our tendon stiffness estimates to previously reported *in vivo* measurements and the model from measures from cadaveric samples, scaled to the *in vivo* geometry of the Achilles tendon (Fig 5b) [4, 8]. The confidence intervals in Fig 5b are one standard deviation of the model estimated by Shaw and Lewis (purple) [4] and measurements from Maganaris and Paul (orange) [8]. The previously reported values of tendon stiffness agree well with each other beyond ~250 N, though the variation is higher in the Shaw and Lewis data [4]. We found that our estimates of tendon stiffness during active muscle contractions are comparable to previously reported values. Outside the lowest levels of force (~<100 N), our estimates of tendon stiffness are within one standard deviation of the model of scaled measurements and previous *in vivo* measurements (Fig 5b) [4, 8].

We also approximated Young’s modulus for comparisons with previous literature. Our approximated Young’s modulus is similar to previously reported values within mammalian tendons at similar levels of stress. Our estimates of Young’s modulus ranged from ~0.03 GPa to ~0.5 GPa, while Young’s modulus from other mammalian tendons ranged from ~0.1 GPa to ~1 GPa at stresses up to 10 MPa [33, 34].

### C. Comparison across days and muscles

Our estimates were repeatable across days even though significant time separated the Day 1 and Day 2 experiments. We used mixed-effects models to evaluate the effect of day on the relationships between ankle stiffness or the static translation ratio and ankle torque. The factor of day was not significant for either dependent variable (Fig 6; ankle stiffness p = 0.54, F_2,3.8_ = 0.73; static translation ratio p = 0.36, F_2,3.8_ = 1.3). The mixed-effects models for ankle stiffness and the static translation ratio fit the data well (ankle stiffness R^2^: 0.95, translation ratio R^2^: 0.93), indicating that the lack of a significant effect was not simply due to high variability across measurements. The mean difference in ankle stiffness and the normalized static translation ratio across days was small. The mean difference in ankle stiffness between Day 1 and Day 2 was 2.4 Nm/rad. This corresponded to an average of 2.7% of the estimated ankle stiffness value for Day 1 across the range of tested torques (0-30 Nm). We observed a mean difference in the normalized static translation ratio across days of −0.014 rad^−1^. The difference across days corresponded to 5.8% of the Day 1 normalized static translation ratio estimates across the range of tested torques.

**Fig 6.**
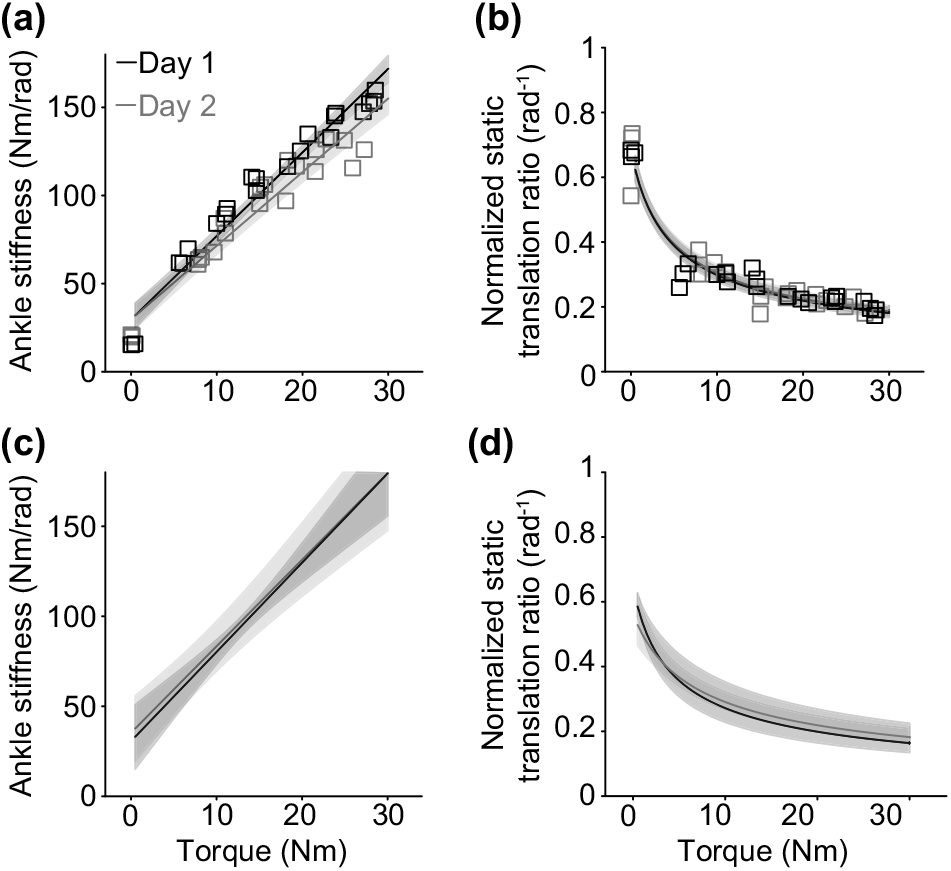
Ankle stiffness (a) and the normalized static translation ratio (b) for a representative subject as a function of plantarflexion torque on Day 1 (black) and Day 2 (grey). Each point represents an individual trial. Group results for ankle stiffness (c) and the normalized static translation ratio (d). The solid line is the predicted ankle stiffness or normalized static translation ratio from the mixed-effects model; the shaded regions are the 95% confidence intervals.

The estimates of ankle stiffness and the normalized static translation ratio did not depend on which MTJ was imaged during conditions with active muscle contractions. Again, we used mixed-effects models as described above but with *muscle* rather than *day* as the additional fixed factor. Data from both days were incorporated into the analysis. Both mixed-effects models fit the data well (stiffness R^2^: 0.93, normalized static translation ratio R^2^: 0.91). Since only one muscle could be imaged at a time, we wanted to ensure we made fair comparisons and that ankle stiffness was matched across muscles. There was no significant difference in the ankle stiffness depending on which muscle was imaged (Fig 7a, c; p = 0.40, F_4,4.1_ = 1.32). Since this result does not depend on which muscle was imaged, it indicates that our experimental conditions were matched across the different imaging sessions.

**Fig 7.**
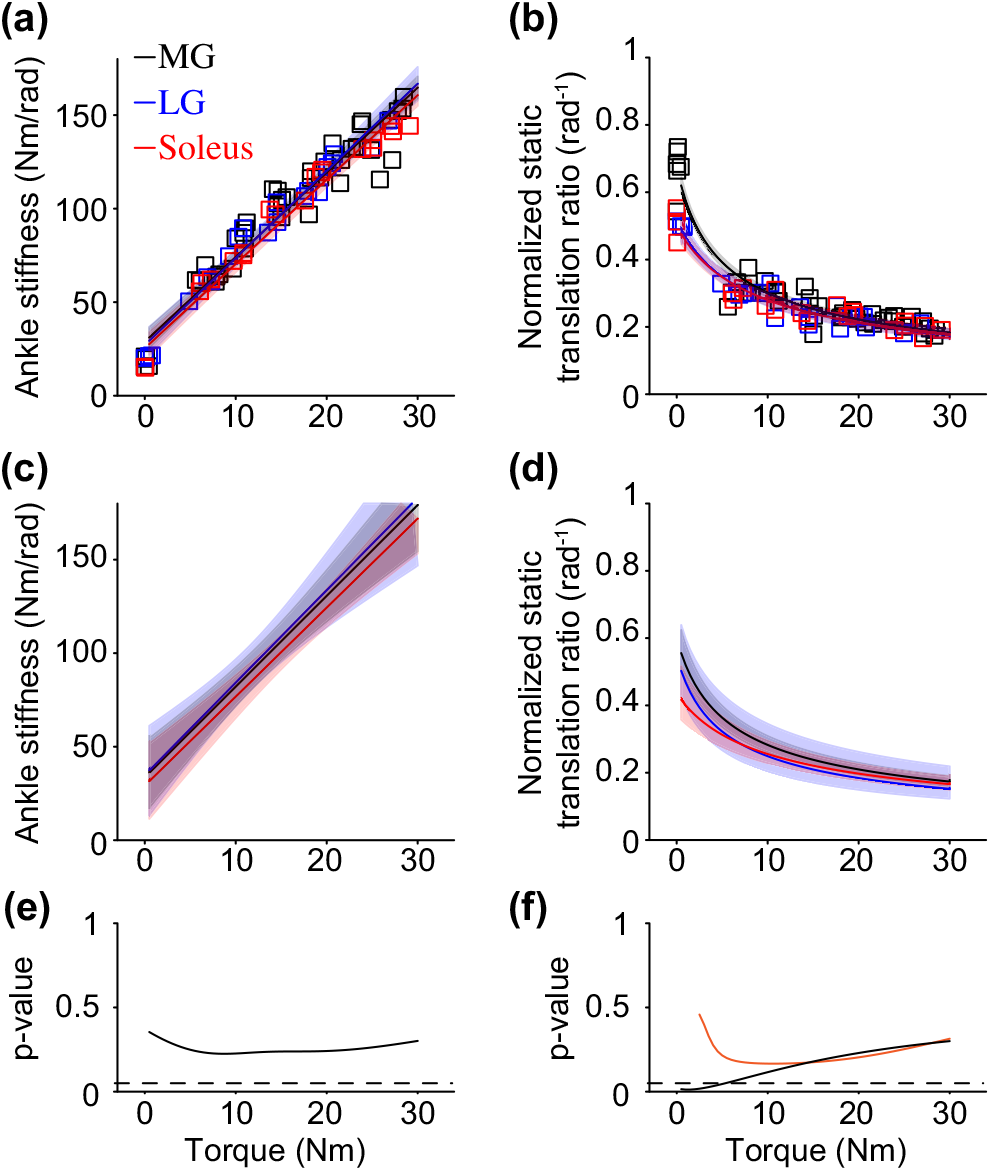
Ankle stiffness (a) and the normalized static translation ratio (b) for a representative subject as a function of the measured plantarflexion torque for the medial gastrocnemius (black), lateral gastrocnemius (blue), and soleus (red). Each point represents an individual trial. Group results for ankle stiffness (c) and the normalized static translation ratio (d). The solid line is the predicted ankle stiffness or normalized static translation ratio from the mixed-effects model; the shaded regions are the 95% confidence intervals. The p-value testing if (e) stiffness or (f) the normalized static translation ratio differed across muscles. The black line is the comparison across muscles on the mixed-effects model that includes the passive data, while the orange line is on the mixed-effects model with only active trials. The dashed line represents p = 0.05.

We did observe a significant difference in the normalized static translation ratio – torque relationship between muscles (Fig 7b, d; p = 0.006, F_4,4.1_ = 20), but this difference was due to measurements made during passive conditions. The post-hoc analysis found significant differences between muscles when the plantarflexion torque was below ~5 Nm (Fig 7f - black line). Across our subjects, we observed a 22 ± 6 %, 29 ± 12%, and 20 ± 33% difference in the normalized static translation ratio during the passive condition between the medial and lateral gastrocnemii, the medial gastrocnemius and soleus, and the lateral gastrocnemius and soleus, respectively. Thus, we tested an additional mixed-effects model that excluded data from passive trials. This model also fit the data well (R^2^ = 0.67) but without any significant differences between muscles (p = 0.26, F_4,4.1_ = 1.96). The orange line illustrates the comparisons when modeling only the active trials in Fig 7f.

### D. Sensitivity Analysis

Our estimates of muscle and tendon stiffness are dependent on knowledge of the Achilles tendon moment arm, which we estimated from the literature. Thus, we used a sensitivity analysis to determine the impact of moment arm accuracy on our estimated stiffnesses across the range of ankle torques tested in this study (Fig 8). Both estimates of muscle and tendon stiffness were sensitive to the accuracy of the moment arm estimate. Over the range of tested conditions, the percentage error in the estimated muscle stiffness would be approximately equal to that in the assumed moment arm value. In contrast, tendon stiffness was much more sensitive to changes in moment arm, especially at low torque levels. On average, the percent error in tendon stiffness was approximately 2.4 times the percent error in the moment arm (Fig 8). Both muscle and tendon stiffness were more sensitive to underestimations (e.g., a negative percent change Fig 8), because they are inversely dependent on the moment arm (Eq 4 & 5).

**Fig 8.**
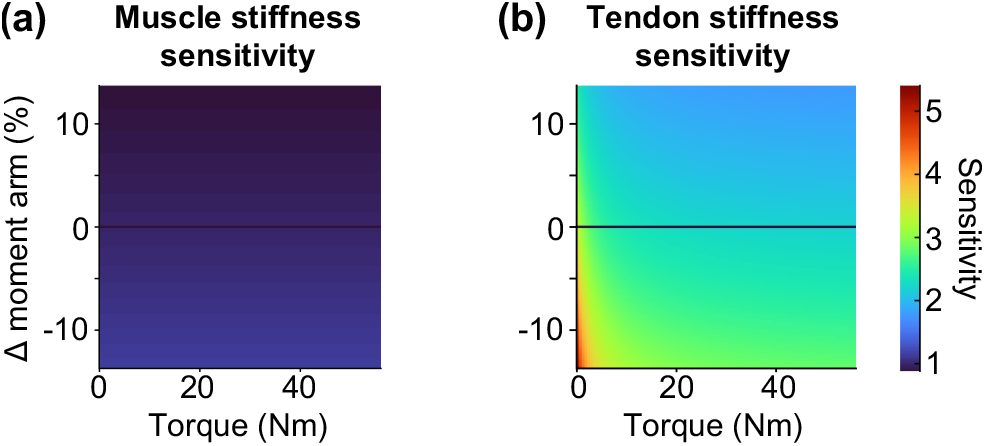
Sensitivity as a function of plantarflexion torque and error in the Achilles tendon moment arm for the measurements of muscle stiffness (a) and tendon stiffness (b). Muscle stiffness measurements were less sensitive to errors in the Achilles tendon moment arm than the measurements of tendon stiffness. Both muscle and tendon stiffness were more sensitive to an underestimation (e.g., negative percent change in the moment arm). Tendon stiffness sensitivity varied with plantarflexion torque, while muscle stiffness sensitivity did not.

## IV. Discussion

We developed a novel *in vivo* technique to quantify joint, muscle, and tendon impedance simultaneously across a range of physiologically relevant activation levels. This was accomplished by combining standard joint-level system identification analysis with B-mode ultrasound imaging. We applied our technique to quantify the impedance of the ankle, the triceps surae muscles, and the Achilles tendon. Our muscle stiffness estimates are consistent with direct measures of muscle short-range stiffness obtained from animal models. Likewise, our tendon stiffness estimates are consistent with previous *in vivo* estimates from human subjects and direct measures from cadaveric samples. Furthermore, the approach yields consistent within-subject estimates even when measurements are separated by long periods of time. Together, our findings demonstrate the validity and reliability of our novel technique for quantifying the mechanical properties of muscle and tendon *in vivo*.

### A. Muscle and tendon stiffness estimates

Our approach is the first that is capable of simultaneous, *in vivo* estimation of muscle and tendon stiffness across a range of activation levels. Our muscle and tendon stiffness estimates are consistent with previously reported measurements (Fig 5) [1, 4, 8]. Our technique is an important innovation for understanding how muscle and tendon stiffness contribute to the net mechanics of the joint. Since both muscle and tendon can be altered by aging and neuromuscular injuries, the ability to identify and track changes in their mechanical properties could be useful for quantifying the effects of aging or injury and the impact of treatments designed to mitigate those effects.

Our novel methodology might be extensible to conditions beyond the isometric situations evaluated in this study. The system identification analyses central to our approach have also been used to estimate joint impedance during movement [20, 35]. Additionally, ultrasound imaging has previously been used to quantify musculotendon motion during movement [36, 37]. Therefore, it is possible that our technique could be applied during similar movement conditions. If so, it would be the first method capable of quantifying muscle and tendon impedance *in vivo* during both posture and movement.

Recently, a theoretical framework has been developed to determine how muscle and tendon stiffness contribute to the net mechanics of the joint [38]. Though the framework has been presented for a similar purpose as our work, its approach to estimation relies on models that may bias the estimated quantities [39]. Such models may be essential when estimating muscle and tendon stiffness during functionally relevant conditions in which the use of perturbation-based system identification is restricted. Our non-parametric technique provides a complementary approach that could be used to validate and refine the models central to the newly proposed framework prior to use in a broader range of conditions.

### B. Methodological considerations

Despite considerable time passing between testing sessions, our estimates were repeatable across days (Fig 6), in accordance with previous measures of ankle impedance [40]. Rather than indicating the reliability of our measure, this finding could suggest that our approach is not sensitive enough to detect changes across time. We do not believe this is the case, as an analysis of each subject, rather than the population, found that one subject had a statistically significant changed ankle stiffness over time and two had significant differences in the normalized static translation ratio. The errors in our population models can be used to estimate the detectable differences of our method across days. We estimate these to be 8% of the average ankle stiffness and 10% of the average normalized static translation ratio – both greater than the average results we observed. These values would, of course, decrease with an increased number of subjects. Our measurements were also insensitive to changes in the ultrasound scanning parameters (frame rate, view area, etc.). This was expected since the Nyquist frequency of the ultrasound scanning rate was well above the frequencies of interest, and our approach to system identification is robust to noise that may result from these minor changes in the image acquisition process. Together, these findings indicate that our method is robust and may be used to track changes in muscle and tendon stiffness that occur over time, especially when muscles are active.

The accuracy of our approach is likely to be compromised in the absence of triceps surae muscle activity. Our analysis assumes that we are quantifying the net impedance of the triceps surae and Achilles tendon. While our normalized static translation ratio—a measure of MTJ displacement relative to ankle displacement—was similar during active conditions regardless of which muscle was imaged, differences were observed between the medial and lateral gastrocnemii and the soleus during passive conditions (Fig 7d). These differences will impact estimates of the net muscle and tendon stiffness, limiting the accuracy of our method during passive conditions. Passive estimates are also likely influenced by our assumption that all plantarflexor torque is transmitted through the Achilles tendon, ignoring contributions from passive structures such as the joint capsule, ligaments, and other musculotendon units, including antagonists, spanning the ankle. These factors can have significant contributions to passive ankle stiffness, but during active conditions, stiffness is much greater [10]. Additionally, nearly all plantarflexor torque about the ankle is produced by the triceps surae [41], which indicates that nearly all the stiffness is from the triceps surae and Achilles tendon. Given these potential sources of error, it is somewhat remarkable that our estimates of muscle and tendon stiffness during passive conditions were still comparable to previously reported values [14, 42, 43]. This may be because the prior studies were susceptible to the same limitations as our method (see *Introduction*), leading to similar errors, or because these limitations were modest at the ankle posture chosen for our measurements.

We found that estimates of tendon stiffness are particularly sensitive to the assumed value for the Achilles tendon moment arm (Fig 8). Both muscle and tendon stiffness estimates were more sensitive to underestimating the moment arm due to how muscle and tendon stiffness are estimated (Eq 4 & 5). When measures of absolute muscle and tendon impedance are critical, accuracy can be increased by making subject-specific measurements of the Achilles tendon moment arm [24, 44, 45]. If only relative measures of muscle and tendon impedance are needed, they can be obtained by estimates of the static translation ratio, which is unaffected by assumed moment arm values.

Given the limited *in vivo* measurements of muscle stiffness, we compared our estimates of muscle stiffness to measurements of short-range stiffness from animals [1], scaled to the human triceps surae. To scale these measurements, we used data from the literature on triceps surae force sharing and fascicle length [2]. We acknowledge that this is only an inference on triceps surae short-range stiffness based on the available information and could be a potential source of error.

Our algorithm for estimating muscle and tendon stiffness assumes that all torque about the ankle is transmitted through the Achilles tendon. This assumption ignores the inertial properties of the foot, which create torques that do not influence muscle or tendon displacement. However, at the frequencies used to estimate stiffness (1 to 3 Hz), frequency response functions have a region of constant magnitude, indicating that inertia had little effect on our estimates of stiffness or the overall impedance within the frequencies relevant to many common tasks such as locomotion (Fig 3 & 4) [46].

## V. Conclusion

We developed an innovative *in vivo* measurement technique to quantify ankle, muscle, and tendon impedance at activation levels relevant to walking and standing. While we applied our approach to the human ankle, it should be feasible to apply this technique to other joints where similar measurements can be made. Our methodology creates new opportunities to investigate the mechanics underlying the control of posture and movement. This ability is critical not only for improving our current understanding of the unimpaired control of posture and movement but also when pathological changes are impacting this control. Our technique may identify the source of the impairment, which can be used for targeted rehabilitation. Insight into the respective roles of the muscle and tendon is also valuable in the design and control of exoskeletons. In this context, the exoskeleton controller could optimize musculotendon mechanics, improving walking efficiency [47].

## References

[1] L. Cui et al., “Modeling short-range stiffness of feline lower hindlimb muscles,” Journal of biomechanics, vol. 41, no. 9, pp. 1945–1952, 2008.

[2] M. Crouzier et al., “Neuromechanical coupling within the human triceps surae and its consequence on individual force-sharing strategies,” Journal of Experimental Biology, vol. 221, no. 21, 2018.

[3] R. E. Kearney and I. W. Hunter, “System identification of human joint dynamics,” Critical reviews in biomedical engineering, vol. 18, no. 1, pp. 55–87, 1990.

[4] K. M. Shaw and G. Lewis, “Tensile properties of human Achilles tendon,” in Proceedings of the 1997 16 Southern Biomedical Engineering Conference, 1997: IEEE, pp. 338–341.

[5] S. Pfeifer et al., “Model-based estimation of knee stiffness,” IEEE Transactions on Biomedical Engineering, vol. 59, no. 9, pp. 2604–2612, 2012.

[6] X. Hu et al., “Muscle short-range stiffness can be used to estimate the endpoint stiffness of the human arm,” Journal of neurophysiology, vol. 105, no. 4, pp. 1633–1641, 2011.

[7] A. D. Vigotsky et al., “Mapping the relationships between joint stiffness, modeled muscle stiffness, and shear wave velocity,” Journal of Applied Physiology, vol. 129, no. 3, pp. 483–491, 2020.

[8] C. N. Maganaris and J. P. Paul, “Tensile properties of the in vivo human gastrocnemius tendon,” J Biomech, vol. 35, no. 12, pp. 1639–46, Dec 2002.

[9] E. J. Rouse et al., “Estimation of human ankle impedance during the stance phase of walking,” IEEE Transactions on Neural Systems and Rehabilitation Engineering, vol. 22, no. 4, pp. 870–878, 2014.

[10] P. Weiss et al., “Human ankle joint stiffness over the full range of muscle activation levels,” Journal of biomechanics, vol. 21, no. 7, pp. 539–544, 1988.

[11] P. Amiri and R. E. Kearney, “Ankle intrinsic stiffness changes with postural sway,” Journal of biomechanics, vol. 85, pp. 50–58, 2019.

[12] J. Trevino and H. Lee, “Sex Differences in 2-DOF Human Ankle Stiffness in Relaxed and Contracted Muscles,” Ann Biomed Eng, vol. 46, no. 12, pp. 2048–2056, Dec 2018.

[13] H. Hauraix et al., “Muscle and tendon stiffness assessment using the alpha method and ultrafast ultrasound,” European journal of applied physiology, vol. 115, no. 7, pp. 1393–1400, 2015.

[14] K. Kubo et al., “Effects of plyometric and isometric training on muscle and tendon stiffness in vivo,” Physiol Rep, vol. 5, no. 15, Aug 2017.

[15] F. E. Zajac, “Muscle and tendon Properties models scaling and application to biomechanics and motor.,” Critical reviews in biomedical engineering, vol. 17, no. 4, pp. 359–411, 1989.

[16] U. Proske and D. Morgan, “Tendon stiffness: methods of measurement and significance for the control of movement. A review,” Journal of biomechanics, vol. 20, no. 1, pp. 75–82, 1987.

[17] K. Nagai et al., “Differences in muscle coactivation during postural control between healthy older and young adults,” Archives of gerontology and geriatrics, vol. 53, no. 3, pp. 338–343, 2011.

[18] I. D. Loram et al., “The passive, human calf muscles in relation to standing: the short range stiffness lies in the contractile component,” The Journal of physiology, vol. 584, no. 2, pp. 677–692, 2007.

[19] E. J. Perreault et al., “Hill muscle model errors during movement are greatest within the physiologically relevant range of motor unit firing rates,” Journal of biomechanics, vol. 36, no. 2, pp. 211–218, 2003.

[20] D. Ludvig et al., “Mechanisms contributing to reduced knee stiffness during movement,” Experimental brain research, vol. 235, no. 10, pp. 2959–2970, 2017.

[21] K. L. Jakubowski et al., “Simultaneous in vivo Estimation of Muscle, Tendon, and Ankle Impedance,” 2020 Annual International Conference of the IEEE Engineering in Medicine and Biology, 2020.

[22] E. J. Perreault et al., “Multiple-input, multiple-output system identification for characterization of limb stiffness dynamics,” Biol Cybern, vol. 80, no. 5, pp. 327–37, May 1999.

[23] A. V. Hill, “The heat of shortening and the dynamic constants of muscle,” Proceedings of the Royal Society of London. Series B-Biological Sciences, vol. 126, no. 843, pp. 136–195, 1938.

[24] E. C. Clarke et al., “A non-invasive, 3D, dynamic MRI method for measuring muscle moment arms in vivo: demonstration in the human ankle joint and Achilles tendon,” Med Eng Phys, vol. 37, no. 1, pp. 93–9, Jan 2015.

[25] F. T. Sheehan, “The 3D in vivo Achilles’ tendon moment arm, quantified during active muscle control and compared across sexes,” J Biomech, vol. 45, no. 2, pp. 225–30, Jan 10 2012.

[26] M. Besomi et al., “Consensus for experimental design in electromyography (CEDE) project: Electrode selection matrix,” J Electromyogr Kinesiol, vol. 48, pp. 128–144, Oct 2019, doi: 10.1016/j.jelekin.2019.07.008.

[27] R. E. Kearney and I. W. Hunter, “Dynamics of human ankle stiffness: variation with displacement amplitude,” J Biomech, vol. 15, no. 10, pp. 753–6, 1982.

[28] D. Miguez et al., “A technical note on variable inter-frame interval as a cause of non-physiological experimental artefacts in ultrasound,” R Soc Open Sci, vol. 4, no. 5, p. 170245, May 2017.

[29] R. F. Kirsch et al., “Muscle stiffness during transient and continuous movements of cat muscle: perturbation characteristics and physiological relevance,” IEEE Transactions on Biomedical Engineering, vol. 41, no. 8, pp. 758–770, 1994.

[30] S. Arya and K. Kulig, “Tendinopathy alters mechanical and material properties of the Achilles tendon,” J Appl Physiol (1985), vol. 108, no. 3, pp. 670–5, Mar 2010.

[31] S. G. Luke, “Evaluating significance in linear mixed-effects models in R,” Behavior research methods, vol. 49, no. 4, pp. 1494–1502, 2017.

[32] I. W. Hunter and R. E. Kearney, “Dynamics of human ankle stiffness: variation with mean ankle torque,” J Biomech, vol. 15, no. 10, pp. 747–52, 1982, doi: 10.1016/0021-9290(82)90089-6.

[33] M. Bennett et al., “Mechanical properties of various mammalian tendons,” Journal of Zoology, vol. 209, no. 4, pp. 537–548, 1986.

[34] L. Cui et al., “In situ estimation of tendon material properties: differences between muscles of the feline hindlimb,” J Biomech, vol. 42, no. 6, pp. 679–85, Apr 16 2009.

[35] H. Lee and N. Hogan, “Time-varying ankle mechanical impedance during human locomotion,” IEEE Transactions on Neural Systems and Rehabilitation Engineering, vol. 23, no. 5, pp. 755–764, 2014.

[36] J. R. Franz et al., “Non-uniform in vivo deformations of the human Achilles tendon during walking,” Gait & posture, vol. 41, no. 1, pp. 192–197, 2015.

[37] G. Lichtwark and A. Wilson, “In vivo mechanical properties of the human Achilles tendon during one-legged hopping,” Journal of Experimental Biology, vol. 208, no. 24, pp. 4715–4725, 2005.

[38] C. P. Cop et al., “Unifying system identification and biomechanical formulations for the estimation of muscle, tendon and joint stiffness during human movement,” Progress in Biomedical Engineering, vol. 3, no. 3, p. 033002, 2021.

[39] E. J. Perreault et al., “Estimation of intrinsic and reflex contributions to muscle dynamics: a modeling study,” IEEE transactions on biomedical engineering, vol. 47, no. 11, pp. 1413–1421, 2000.

[40] R. E. Kearney et al., “System identification of human ankle dynamics: intersubject variability and intrasubject reliability,” Clin Biomech (Bristol, Avon), vol. 5, no. 4, pp. 205–17, Nov 1990, doi: 10.1016/0268-0033(90)90004-P.

[41] S. R. Ward et al., “Are current measurements of lower extremity muscle architecture accurate?,” Clinical orthopaedics and related research, vol. 467, no. 4, pp. 1074–1082, 2009.

[42] A. Konrad et al., “Acute effects of constant torque and constant angle stretching on the muscle and tendon tissue properties,” Eur J Appl Physiol, vol. 117, no. 8, pp. 1649–1656, Aug 2017.

[43] A. Konrad and M. Tilp, “Increased range of motion after static stretching is not due to changes in muscle and tendon structures,” Clinical Biomechanics, vol. 29, no. 6, pp. 636–42, Jun 2014.

[44] K. Manal et al., “A hybrid method for computing achilles tendon moment arm using ultrasound and motion analysis,” Journal of applied biomechanics, vol. 26, no. 2, pp. 224–228, 2010.

[45] J. R. Franz et al., “Ankle Rotation and Muscle Loading Effects on the Calcaneal Tendon Moment Arm: An In Vivo Imaging and Modeling Study,” Annals of biomedical engineering, vol. 47, no. 2, pp. 590–600, 2019.

[46] C. Angeloni et al., “Frequency content of whole body gait kinematic data,” IEEE Transactions on rehabilitation engineering, vol. 2, no. 1, pp. 40–46, 1994.

[47] R. W. Nuckols et al., “Ultrasound imaging links soleus muscle neuromechanics and energetics during human walking with elastic ankle exoskeletons,” Scientific reports, vol. 10, no. 1, pp. 1–15, 2020.

