## Supplemental Material for "Simultaneous quantification of ankle, muscle, and tendon impedance in humans"

In this study, we used mixed-effects models to determine if ankle and the normalized static translation ratio differed across days and when imaging different muscles. To ensure transparency and repeatability of our analyses, we present the mixed-effects models used in this study.

#### *A. Effect of day on ankle stiffness*

For this analysis, we used a linear mixed-effects model with subject treated as a random factor, plantarflexion torque as a continuous factor, and day as a fixed factor, with an interaction term between day and torque. Effects coding was used for all analyses. Ankle stiffness ($K_A$) was fit as a function of plantarflexion torque as follows:

$$K_A = \beta_0 + \beta_{0Day} + \beta_1 \cdot Torque + \beta_{1Day} \cdot Torque + b_{0i} + b_{1i} \cdot Torque + b_{0iDay} + b_{1iDay} \cdot Torque + \varepsilon \quad \text{(S1)}$$

$$[b_{0i}\ b_{1i}\ b_{0iDay}\ b_{1iDay}] \sim N(0, \Psi), \varepsilon \sim N(0, \sigma^2) \quad \text{(S2)}$$

Where $\beta_0$ and $\beta_1$ are the intercept and slope across all subjects, respectively, $\beta_{0Day}$ and $\beta_{1Day}$ are the differences in the intercept and slope across days for all subjects. $b_{0i}$, $b_{1i}$, $b_{0iDay}$, and $b_{1iDay}$ are the intercept and slope for the $i^{th}$ subject, including an interaction with day. $b_{0i}$, $b_{1i}$, $b_{0iDay}$, and $b_{1iDay}$ come from a normal distribution with the covariance ($\Psi$) shown in Table S3. We have also included the fixed-effect estimates (Table S1) and the fixed-effect covariance matrix (Table S2).

Supplemental Table I

Fixed effects for the effect of torque and day on ankle stiffness

| *Parameter* | *Estimate* |
|---|---|
| $\beta_0$ | 32.9 Nm/rad |
| $\beta_{0Day}$ | 2.3 Nm/rad |
| $\beta_1$ | 4.9 Nm/rad/Nm |
| $\beta_{1Day}$ | -0.07 Nm/rad/Nm |

Supplemental Table 2

Variance-covariance matrix for the fixed effects parameters shown in supplemental table 1

| | $\beta_0$ | $\beta_{0Day}$ | $\beta_1$ | $\beta_{1Day}$ |
|---|---|---|---|---|
| $\beta_0$ | 83 | 0.1 | -5 | 0.5 |
| $\beta_{0Day}$ | 0.1 | 4 | 0.4 | -0.01 |
| $\beta_1$ | -5 | 0.4 | 0.5 | 0.003 |
| $\beta_{1Day}$ | 0.5 | -0.01 | 0.003 | 0.02 |

Supplemental Table 3

Variance-covariance estimates for the random effect show in eq s2

| *Parameter* | *Estimate* |
|---|---|
| $\Psi$ | $\begin{bmatrix} 240 & -1 & -15 & 2 \\ -1 & 5 & 1 & 0.4 \\ -15 & 1 & 1 & -0.01 \\ 2 & 0.4 & -0.01 & 0.04 \end{bmatrix}$ |
| $\sigma^2$ | 131 |

#### *B. Effect of day on the normalized static translation ratio*

For this analysis, we used a generalized linear mixed-effects model with subject treated as a random factor, plantarflexion torque as a continuous factor, and day as a fixed factor, with an interaction term between day and torque. The normalized static translation ratio ($\tilde{T}_R$) was fit as a function of plantarflexion torque as follows:

$$\tilde{T}_R = g^{1/p} \quad \text{(S3)}$$

$$g = \beta_0 + \beta_{0Day} + \beta_1 \cdot Torque + \beta_{1Day} \cdot Torque + b_{0i} + b_{1i} \cdot Torque + b_{0iDay} + b_{1iDay} \cdot Torque + \varepsilon \quad \text{(S4)}$$

$$[b_{0i}\ b_{1i}\ b_{0iDay}\ b_{1iDay}] \sim N(0, \Psi), \varepsilon \sim N(0, \sigma^2) \quad \text{(S5)}$$

We used a power link function of $p$ = -1.9, and $g$ is the linear prediction of the fixed and random effects for the generalized linear model. Therefore, the variables are the same as Section A.

Supplemental Table 4

Fixed effects for the effect of torque and day on the normalized static translation ratio

| *Parameter* | *Estimate* |
|---|---|
| $\beta_0$ | 2.6 rad$^{-1}$ |
| $\beta_{0Day}$ | -0.4 rad$^{-1}$ |
| $\beta_1$ | 0.9 rad$^{-1}$ Nm$^{-1}$ |
| $\beta_{1Day}$ | 0.1 rad$^{-1}$ Nm$^{-1}$ |

Supplemental Table 5

Variance-covariance matrix for the fixed effects parameters shown in supplemental table 4

| | $\beta_0$ | $\beta_{0Day}$ | $\beta_1$ | $\beta_{1Day}$ |
|---|---|---|---|---|
| $\beta_0$ | 0.08 | -0.06 | 0.01 | 0.02 |
| $\beta_{0Day}$ | -0.06 | 0.05 | -0.01 | -0.03 |
| $\beta_1$ | 0.01 | -0.01 | 0.006 | 0.01 |
| $\beta_{1Day}$ | 0.02 | -0.03 | 0.01 | 0.03 |

Supplemental Table 6

Variance-covariance estimates for the random effect show in eq s5:

| *Parameter* | *Estimate* |
|---|---|
| $\Psi$ | $\begin{bmatrix} 0.2 & -0.2 & 0.04 & 0.06 \\ -0.2 & 0.1 & 0.04 & -0.08 \\ 0.04 & -0.04 & 0.02 & 0.03 \\ 0.06 & -0.08 & 0.03 & 0.08 \end{bmatrix}$ |
| $\sigma^2$ | 0.002 |

#### *C. Effect of muscle on ankle stiffness*

For this analysis, we used a linear mixed-effects model with subject treated as a random factor, plantarflexion torque as a continuous factor, muscle as a fixed factor with an interaction term between muscle and torque. Effects coding was used for all analyses. Ankle stiffness ($K_A$) was fit as a function of plantarflexion torque as follows:

$$K_A = \beta_0 + \beta_{0Musclei} + \beta_1 \cdot Torque + \beta_{1Musclei} \cdot Torque + b_{0i} + b_{1i} \cdot Torque + b_{0iMusclei} + b_{1iMusclei} \cdot Torque + \varepsilon \quad \text{(S6)}$$

$$[b_{0i}\ b_{1i}\ b_{0iMusclei}\ b_{1iMusclei}] \sim N(0, \Psi), \varepsilon \sim N(0, \sigma^2) \quad \text{(S7)}$$

Where $\beta_0$ and $\beta_1$ are the intercept and slope across all subjects, respectively, $\beta_{0Muscle}$ and $\beta_{1Muscle}$ are the differences in the intercept and slope across muscles for all subjects. $b_{0i}$, $b_{1i}$, $b_{0iMuscle}$, and $b_{1iMuscle}$ are the intercept and slope for the $i^{th}$ subject, including an interaction with muscle. $b_{0i}$, $b_{1i}$, $b_{0iMuscle}$, and $b_{1Muscle}$ come from a normal distribution with the covariance ($\Psi$) shown in Table S9. We have also included the fixed-effect estimates (Table S7) and the covariance matrix for the fixed-effects (Table S8).

SUPPLEMENTAL TABLE 7
FIXED EFFECTS FOR THE EFFECT OF TORQUE AND MUSCLE ON ANKLE STIFFNESS

| *Parameter* | *Estimate* |
|---|---|
| $\beta_0$ | 32.5 Nm/rad |
| $\beta_{0Muscle1}$ | 1.0 Nm/rad |
| $\beta_{0Muscle2}$ | 2.3 Nm/rad |
| $\beta_1$ | 4.9 Nm/rad/Nm |
| $\beta_{1Muscle1}$ | 0.003 Nm/rad/Nm |
| $\beta_{1Muscle2}$ | 0.09 Nm/rad/Nm |

SUPPLEMENTAL TABLE 8
VARIANCE-COVARIANCE MATRIX FOR THE FIXED EFFECTS PARAMETERS SHOWN IN SUPPLEMENTAL TABLE 7

| | $\beta_0$ | $\beta_{0Muscle1}$ | $\beta_{0Muscle2}$ | $\beta_1$ | $\beta_{1Muscle1}$ | $\beta_{1Muscle2}$ |
|---|---|---|---|---|---|---|
| $\beta_0$ | 115 | -12 | 17 | -7 | 0.8 | -2 |
| $\beta_{0Muscle1}$ | -12 | 4 | -3 | 0.7 | -0.2 | 0.3 |
| $\beta_{0Muscle2}$ | 17 | -3 | 7 | -0.8 | 0.2 | -0.4 |
| $\beta_1$ | -7 | 0.7 | -0.8 | 0.5 | -0.05 | 0.2 |
| $\beta_{1Muscle1}$ | 0.8 | -0.2 | 0.2 | -0.05 | 0.01 | -0.02 |
| $\beta_{1Muscle2}$ | -2 | 0.3 | -0.4 | 0.2 | -0.02 | 0.06 |

SUPPLEMENTAL TABLE 9
VARIANCE-COVARIANCE ESTIMATES FOR THE RANDOM EFFECT SHOW IN EQ S7

| *Parameter* | *Estimate* |
|---|---|
| $\Psi$ | $\begin{bmatrix} 338 & -35 & 50 & -20 & 2 & -7 \\ -35 & 4 & -5 & 2 & -0.3 & 0.7 \\ 50 & -5 & 9 & -2 & 0.4 & -0.8 \\ -20 & 2 & -2 & 2 & -0.1 & 0.4 \\ 2 & -0.3 & 0.4 & -0.1 & 0.02 & -0.05 \\ -7 & 0.7 & -0.8 & 0.4 & -0.05 & 0.1 \end{bmatrix}$ |
| $\sigma^2$ | 166 |

### *D. Effect of muscle on the normalized static translation ratio*

For this analysis, we used a generalized linear mixed-effects model with subject treated as a random factor, plantarflexion torque as a continuous factor, muscle as a fixed factor with an interaction term between muscle and torque. The normalized static translation ratio ($\tilde{T}_R$) was fit as a function of plantarflexion torque as follows:

$$\tilde{T}_R = g^{1/p} \quad (S8)$$

$$g = \beta_0 + \beta_{0Musclei} + \beta_1 \cdot Torque + \beta_{1Musclei} \cdot Torque + b_{0i} + b_{1i} \cdot Torque + b_{0iMusclei} + b_{1iMusclei} \cdot Torque + \varepsilon \quad (S9)$$

$$[b_{0i}\ b_{1i}\ b_{0iMusclei}\ b_{1iMusclei}] \sim N(0, \Psi), \varepsilon \sim N(0, \sigma^2) \quad (S10)$$

We used a power link function of p = -1.9, and *g* is the linear prediction of the fixed and random effects for the generalized linear model. Therefore, the variables are the same as Section C.

SUPPLEMENTAL TABLE 10
FIXED EFFECTS FOR THE EFFECT OF TORQUE AND MUSCLE ON THE NORMALIZED STATIC TRANSLATION RATIO

| *Parameter* | *Estimate* |
|---|---|
| $\beta_0$ | 3.5 $rad^{-1}$ |
| $\beta_{0Muscle1}$ | -0.9 $rad^{-1}$ |
| $\beta_{0Muscle2}$ | -0.4 $rad^{-1}$ |
| $\beta_1$ | 0.9 $rad^{-1}\,Nm^{-1}$ |
| $\beta_{1Muscle1}$ | -0.08 $rad^{-1}\,Nm^{-1}$ |
| $\beta_{1Muscle2}$ | 0.2 $rad^{-1}\,Nm^{-1}$ |

SUPPLEMENTAL TABLE 11
VARIANCE-COVARIANCE MATRIX FOR THE FIXED EFFECTS PARAMETERS SHOWN IN SUPPLEMENTAL TABLE 10

| | $\beta_0$ | $\beta_{0Muscle1}$ | $\beta_{0Muscle2}$ | $\beta_1$ | $\beta_{1Muscle1}$ | $\beta_{1Muscle2}$ |
|---|---|---|---|---|---|---|
| $\beta_0$ | 0.07 | -0.002 | -0.09 | 0.02 | -0.009 | 0.02 |
| $\beta_{0Muscle1}$ | -0.002 | 0.02 | 0.01 | 0.007 | -0.004 | 0.01 |
| $\beta_{0Muscle2}$ | -0.09 | 0.01 | 0.5 | 0.02 | -0.003 | 0.06 |
| $\beta_1$ | 0.02 | 0.007 | 0.02 | 0.01 | -0.007 | 0.02 |
| $\beta_{1Muscle1}$ | -0.009 | -0.004 | -0.003 | -0.007 | 0.004 | -0.009 |
| $\beta_{1Muscle2}$ | 0.02 | 0.01 | 0.06 | 0.02 | -0.009 | 0.03 |

SUPPLEMENTAL TABLE 12
VARIANCE-COVARIANCE ESTIMATES FOR THE RANDOM EFFECT SHOW IN EQ S10

| *Parameter* | *Estimate* |
|---|---|
| $\Psi$ | $\begin{bmatrix} 0.2 & 0.02 & -0.3 & 0.05 & -0.03 & 0.05 \\ 0.02 & 0.009 & 0.03 & 0.02 & -0.009 & 0.03 \\ -0.3 & 0.03 & 1 & 0.05 & -0.007 & 0.2 \\ 0.05 & 0.02 & 0.05 & 0.04 & -0.02 & 0.06 \\ -0.03 & -0.009 & -0.007 & -0.01 & 0.009 & -0.02 \\ 0.05 & 0.03 & 0.2 & 0.06 & -0.02 & 0.09 \end{bmatrix}$ |
| $\sigma^2$ | 0.002 |
